# A trade-off between desiccation resistance and social communication in a social bee

**DOI:** 10.64898/2026.11.19.767937

**Authors:** Seyed Ali Modarres Hasani, Nathan Derstine, Etya Amsalem

## Abstract

The cuticular lipids of insects serve both as a barrier against desiccation and as chemical signals. In social insects, these lipids regulate reproduction and nestmate recognition, potentially creating a trade-off between waterproofing and communication. We tested this hypothesis in bumblebee workers by examining the roles of cuticular compounds in desiccation resistance and behavioral attraction. Desiccation survival varied with age, social state, and ovary activation, with fertile queenless workers showing the greatest resistance. Cuticular profiles changed markedly with age and social environment. Although the total amount of secretion remained similar among groups, queenright workers showed age-dependent decline in alkenes and a corresponding increase in two acetate esters (tetracosyl and hexacosyl acetate). To determine the functions of these compounds, we tested their effects on desiccation survival and attraction. Topical application of several hydrocarbons, but not tetracosyl acetate, increased desiccation resistance. In contrast, tetracosyl acetate, but not hydrocarbons, elicited attraction, and only in queenright workers. These results demonstrate an age-dependent shift in cuticular lipid composition in the presence of the queen from producing waterproofing hydrocarbons toward signaling compounds in the presence of the queen, reducing desiccation resistance while enhancing communication. Our findings suggest that the evolution of insect cuticular lipids has been shaped by both physiological demands and selection for complex social communication.

**Summary statement:** Bumblebees balance waterproofing and social communication by shifting their surface lipids, revealing how social life can reshape traits that are essential for coping with environmental stress.

## Background

The insect cuticle is coated with a complex mixture of lipids that plays a central role in mediating interactions between the organism and its environment (Blomquist et al., 2018, Howard and Blomquist, 2005, Blomquist and Ginzel, 2021). These surface lipids, dominated by long chain saturated and unsaturated hydrocarbons along with smaller amounts of esters, fatty acids, alcohols, and other derivatives, form the primary barrier limiting water loss and protecting insects from desiccation and microorganism activity (Gibbs, 1998, Wrońska et al., 2023). At the same time, cuticular compounds function as chemical signals that convey information such as species identity, age, health, reproductive status, and social context (Barbero, 2016, Smith and Liebig, 2017, Blomquist and Ginzel, 2021). Because the same classes of compounds contribute both to waterproofing and to communication, the composition of cuticular lipids is expected to reflect a balance between physiological demands and signaling functions. These roles can impose conflicting selective pressures: traits that maximize barrier efficiency may not optimize signal transmission or perception (Berson et al., 2019). Understanding how insects resolve this potential trade-off is therefore central to explaining the evolution and plasticity of cuticular lipid profiles.

Avoiding desiccation is a challenging task for terrestrial insects because of their small body size and large surface area-to-volume ratio (Chown et al., 2011, Gibbs, 2002). To reduce water permeability through the cuticle, insects produce cuticular lipids that can be modified in various ways as insects age, reproduce or in response to environmental changes (Menzel et al., 2017, Blomquist and Ginzel, 2021). Aging, for example, may lead to reduction in the amount of long-chain hydrocarbons, making species such as mosquitoes and flies more sensitive to desiccation (Lyons et al., 2014, Kárpáti et al., 2023). Likewise, reproductive maturation is often associated with changes in insect cuticular composition, linked to mate attraction and signaling reproductive status (Blomquist and Bagnères, 2010, Ferveur, 2005). Cuticular lipids may also change in response to temperature or elevation. Desert beetles (*Eleodes armata* and termite nymphs (*Cryptotermes brevis)*, for example, both produce higher quantities of hydrocarbons and a greater proportion of long-chain components as temperatures increase (Hadley, 1977, Woodrow et al., 2000). *Drosophila melanogaster* shift from producing short to long-chain cuticular hydrocarbons (Rajpurohit et al., 2021) whereas ants reduce the proportion of unsaturated and methyl branched alkanes as temperatures increase (Menzel et al., 2018). Changes in chain length or in the abundance of straight and methyl-branched alkanes in response to elevation have been further reported in termites (Camarota et al., 2025), fruit flies (Nair et al., 2025) and bumblebees (Narah et al., 2024).

Cuticular lipid profiles are species-specific and have been co-opted as chemical signals in both solitary and social insects. Across species, they convey information on population and species identity, sex, age, nest, mating and reproductive status, and mate quality (Chung and Carroll, 2015, Leonhardt et al., 2016, Blomquist and Ginzel, 2021). In social species, they have further evolved to mediate nest identity and caste recognition (Smith and Liebig, 2017). For instance, cuticular compounds such as n-alkanes and (Z)-9-alkenes function as key nestmate recognition signals across multiple *Formica* ant species (Akino et al., 2004, Martin et al., 2008). Cuticular lipids further play an important role in conveying information about fecundity with fertile females often producing different composition and quantity of cuticular lipids (Amsalem, 2020). Bumblebee queens, for example, have three times more total hydrocarbons on their cuticle than non-reproductive queens alongside a higher short-to-long chain ratio (Orlova et al., 2020). In some cases, cuticular hydrocarbons were shown to inhibit reproduction or reproductive-related behaviors in subordinate workers (Holman et al., 2010, Funaro et al., 2018, Van Oystaeyen et al., 2014). In ants, selected compounds signal dominance via specific cuticular alkenes such as (*Z*)-9-nonacosene in *Odontomachus brunneus* (Smith et al., 2012) and 9-hentriacontene in *Dinoponera quadriceps*(Peeters et al., 1999) or through overall chain elongation and the presence of 13,23-dimethylheptatriacontane in *Harpegnathos saltator* (Liebig et al., 2000).

The dual function of cuticular lipids, particularly in social species, raises the question of whether their signaling role comes at a cost. Signal specificity and reliability are often achieved through greater chemical diversity and chemical modifications which may contrast with the requirements of an effective barriers against water loss (Chung and Carroll, 2015, Hefetz, 2026). Resistance to desiccation is generally enhanced by cuticular blends with high melting temperatures that pack tightly and form stable waterproofing layers. Accordingly, chemical blends composed of long chain, saturated, and low-volatility compounds are expected to reduce water permeability. In contrast, signals often rely on chemically diverse blends that increase specificity and information content, including unsaturated, branched, or otherwise modified compounds that reduce packing efficiency and melting temperature. In addition, effective chemical communication requires compounds to be detectable by receivers, favoring greater chemical distinctiveness and, in some cases, increased diffusibility. Thus, traits that improve signaling may compromise waterproofing properties, whereas traits that maximize desiccation resistance may reduce signal complexity and detectability (Leonhardt et al., 2016, Perez and Aron, 2020, Hefetz, 2026).

Such a trade-off between signaling and protective functions has been suggested in several studies, particularly in ants. For example, (Menzel et al., 2019) proposed that branched and chemically-diverse hydrocarbons may improve signal specificity and information content, but at the same time reduce the efficiency of the cuticle as a waterproofing barrier. A detailed example from Argentine ants demonstrate they exhibit reduced levels of complex trimethyl alkanes under severe desiccation pressure (Buellesbach et al., 2018). Because trimethyl alkanes are structurally diverse compounds thought to contribute strongly to colony recognition and signaling specificity, their reduction under dry conditions may reflect a shift away from communication and toward waterproofing. Similarly, a study on two sister species of *Lasius* ants showed that dimethyl and trimethyl alkanes contributed more strongly to colony differences, whereas alkanes and monomethyl alkanes were more strongly associated with acclimatory responses (Wittke et al., 2022). In *Myrmica* ants, increased production of cuticular hydrocarbons is associated with reduced survival under drought conditions, likely reflecting selection for signaling at the expense of waterproofing (Sprenger et al., 2018). Together, these studies suggest that compounds involved in social communication may be under selective pressures that differ from, and potentially conflict with, those optimizing desiccation resistance. However, direct empirical evidence for this trade-off, particularly under natural conditions that do not involve extreme environmental changes and outside of ants, remains limited.

In this study, we examine the trade-off between the functional role of cuticular lipids in waterproofing and their signaling role in the bumblebee *Bombus impatiens*. This species is a eusocial annual bee in which founder queens must survive extended winter diapause, and colonies are relatively small and typically initiated underground. In newly founded nests, workers act as helpers whereas the queen is the sole reproductive female (Amsalem et al., 2015). However, as colonies and workers age, or in the absence of the queen, workers engage in reproductive competition with the queen and among themselves and attempt to lay unfertilized eggs that develop into males. These shifts in worker reproductive physiology and social organization are accompanied by changes in the composition of their chemical secretion, with demonstrated differences between queenright (QR) and queenless (QL) workers (i.e., in the presence and absence of the queen), workers of different age, and workers from young colonies vs. colonies that rear sexuals (Derstine et al., 2023, Orlova et al., 2022, Derstine et al., 2021, Amsalem et al., 2014a, Amsalem et al., 2013b, Amsalem and Hefetz, 2011, Amsalem et al., 2009). The complex sociobiology and ecology of bumblebees highlight the dual importance of maintaining an effective barrier against water loss while preserving the chemical communication essential for social organization.

We first examined the desiccation resistance of workers and how it is affected by their social state, age, reproductive physiology and mass. To link these traits with the cuticular water barrier function, we analyzed the cuticular lipid secretion of workers subjected to the same social and age treatments. We then examined the role of selected cuticular compounds in desiccation stress and in communication by testing worker survival following a topical application or by examining workers attraction to the same compounds. We hypothesized that the primary function of worker cuticular lipids is to prevent desiccation. However, because both the total amount and composition of these secretions are constrained, stage-dependent investment in compounds that serve in social communication is expected to come at a cost to desiccation resistance.

## Methods

### Bees

Colonies of *B. impatiens*, each containing a queen, brood, and workers, were obtained from Koppert Biological Systems (Howell, MI, USA). Colonies were maintained in nest boxes with transparent top lids within a dark environmental chamber maintained at a temperature of 28-30 °C and 60% relative humidity. Bees were fed *ad libitum* with a 60% sucrose solution and fresh, honeybee-collected pollen. To obtain workers of known age and social state, we collected newly-emerged workers (“callows”, identified by their silvery-grey appearance) and assigned them to one of two social treatments: queenright (QR) or queenless (QL). Bees were collected progressively as they emerged in their maternal colonies, and assignment to treatments within each experiment was conducted in parallel. QR workers were individually marked with a number tag and returned to their maternal colonies. QL workers were removed from their maternal colonies upon emergence and placed in plastic cages (11 cm diameter × 7 cm height) in groups of three. Workers were sampled upon reaching the target age (2,6, or 10 days). Cages were kept under the same conditions as the maternal colonies and were provided with the same food. In total, we used 25 colonies, from which we sampled 667 workers for the various analyses (see Table S1 for detailed sample sizes).

### Desiccation survival assays

To assess desiccation resistance, individual workers were placed in a glass apparatus (Wang et al., 2022) within an incubator maintained at 35 °C and 10% relative humidity. We used these conditions to impose strong desiccation stress, as high temperatures accelerate water loss and 35°C represents a biologically relevant thermal challenge for *B. impatiens* (Quinlan et al., 2023, Bretzlaff et al., 2023, Botsch et al., 2024). The apparatus consisted of two 50 mL glass tubes (24 mm diameter × 94 mm length) connected and separated by a thin mesh. One arm contained 5 g of silica gel, while the other housed the bee (Figure S1). Bees were monitored hourly until death.

We examined the survival of QR and QL workers at ages 2, 6, and 10 days (11-15 bees per treatment), in addition to 15 callow bees. Body mass was measured at the onset of the experiment and upon death. At the end of the experiment, workers were dissected to assess ovarian activation by measuring the average length of three of the terminal oocytes as in (Amsalem et al., 2009).

### Chemical analysis of worker cuticular lipids

A second set of worker samples of the same ages and social states was collected, frozen immediately and stored at −80 °C until analysis (11–15 bees per treatment). Workers were individually extracted in 1mL hexane for 1 minute. The extract was passively evaporated to approximately 150 µL and then transferred to a glass insert within a 2 mL glass vial. The solvent was nearly completely evaporated, and the extract was resuspended in hexane containing 5 µg of eicosane (C20) as an internal standard (1 µg/µL).

Samples were analyzed on an Agilent 7890A GC equipped with an HP-5ms column (0.25 mm ID × 30 m × 0.25 µm film thickness, Agilent Technologies, Santa Clara, CA) coupled to an Agilent 5975C mass spectrometer. Runs were performed in split less mode with a temperature program starting at 60 °C, increasing at 15 °C/min to 120 °C, then at 4 °C/min to 300 °C, with a final hold of 5 min. Helium was used as the carrier gas, with an injection volume of 1 µL. The injector and FID temperatures were maintained at 250°C and 320°C, respectively. Chromatograms and spectra were analyzed using MSD ChemStation software (Agilent). Peaks were identified using the NIST database and by comparing retention times and mass spectra with synthetic compound standards.

All samples were quantified using a Trace 1310 gas chromatograph (Thermo Scientific) equipped with a flame-ionization detector (FID) and a TG-5MS column (0.25 mm id x 30 m x 0.25 μm film thickness, Thermo Fisher). Runs were performed in split less mode using the same conditions and temperature program as above. Chromatograms were integrated using Chromeleon 7.0 software (Thermo Scientific). Compounds were identified based on diagnostic ions in the mass spectra and by matching retention times and spectra with authentic standards. Peak areas were normalized to the internal standard and are presented as either absolute quantities or percentages of the total secretion.

### Desiccation survival assays

To directly test the role of specific cuticular lipids in waterproofing, we measured worker survival under desiccation following a topical application of selected compounds. In the first round, we tested 10 of the most abundant compounds in the cuticular lipids of workers, based on the SIMPER analysis. These included 6 alkanes (C23, C24, C25, C27, C29, C31), 3 alkenes (C23:1(Z7), C25:1 (Z7), C25:1 (Z9)) and one acetate ester (tetracosyl acetate, TCA). Compounds were applied in a single dose of 50 µg within a volume of 1 µl to the thorax of callow bees prior to placing them in the glass apparatus (11-15 bees/treatment). All compounds were dissolved in hexane, and control bees received an application of 1 µl of hexane. In the second round, we repeated this experiment with 6 compounds (C23:1 (Z7), C25:1 (Z7), C27, C29, C31 and TCA), and applied them to the thorax of QL, 10 d old bees in a single dose of 400 µg within the same volume.

### Attraction bioassays

Behavioral assays were conducted using a T maze (Figure S1) (Amsalem and Hefetz, 2026). We tested the attraction of workers from two social states, QR and QL, to three selected compounds, an alkane (C27), an alkene (C25:1 (Z7), and an acetate ester (TCA). Overall, we conducted 168 individual bioassays, 13-16 bioassays per compound. QR workers were sampled directly from two commercial colonies upon arrival for 3-5 consecutive days, whereas QL workers were taken from the same colonies and placed in small cages of three for 3-5 days before their behavior was examined. Every worker was tested only once. The compounds were selected based on the outcome of the previous, their molecular weight and volatility to ensure that their physical properties were as similar as possible. C27 has a molecular weight of 380.7 g/mol and an estimated vapor pressure of about 10⁻⁸ to 10⁻⁹ Pa. C25:1 (7Z) has a molecular weight of 350.7 g/mol and an estimated vapor pressure of about 10⁻⁶ to 10⁻⁷ Pa. TCA has a molecular weight of 396.7 g/mol and an estimated vapor pressure of about 10⁻⁷ to 10⁻⁸ Pa. Overall, all three compounds have low and comparable volatility.

The T maze was made of polycarbonate sheets (Figure S2) (Amsalem and Hefetz, 2026). The endpoint of each arm contained a cotton wick with either hexane (solvent control) or a treatment odor at a single dose of 400 µg in a volume of 1 µl, the same concentration applied to the bee cuticle in the survival assays. Light was blocked from both sides of the maze using dark sheets. Tests were conducted in the morning under ambient lighting conditions in a normally lit laboratory. The solvent was allowed to evaporate from the cotton wicks for 1 minute before they were placed in the maze. Each arm led to a plastic cage. Each worker was given 10 minutes to choose an arm. A choice was scored as positive when the worker passed the cotton wick and entered the cage without the possibility of returning to the maze. Workers that did not choose within 10 minutes were classified as non responsive. Odor positions were alternated between arms and workers showed no preference to either of the sides. The maze was wiped cleaned between trials using ethanol. All bioassays were conducted during the morning hours in ambient light. Dark cardboard sheets were placed on both sides of the maze to ensure similar light levels in both arms.

All alkanes and alkenes were purchased from Sigma. Tetracosyl acetate (TCA) was synthesized in the lab by acetylation of tetracosanol following a standard esterification procedure. Tetracosanol (240 mg) was dissolved in pyridine (3 mL), and 4-dimethylaminopyridine (DMAP; 80 mg) was added as a catalyst. Acetic anhydride was added in excess (≥2 equivalents), and the reaction mixture was stirred at room temperature overnight to ensure complete conversion of the alcohol to the acetate ester. The reaction was then quenched with ethanol (5 equivalents relative to acetic anhydride) and stirred for several hours to consume residual acetic anhydride. The reaction mixture was diluted with water and extracted with hexane. The organic phase was washed with brine, dried over anhydrous sodium sulfate and concentrated by passive evaporation. This crude product was purified by vacuum flash chromatography followed by recrystallization from hexane. The resulting product formed white crystalline solids consistent with tetracosyl acetate, as was confirmed by GC/MS analysis which matched published spectra and retention indices.

### Data Analysis

Data were analyzed using JMP Pro 18 and RStudio. Time to death (h) was analyzed using a linear model with social condition, age, and their interaction as fixed effects to account for all variables within a single model. Because all individuals were monitored until death, no observations were censored, and survival time was treated as a continuous response variable. The average terminal oocyte size and mass were analyzed in the same way. Correlations between time to death and oocyte size, initial mass, and mass loss were calculated using Pearson correlation.

Cuticular compound composition was evaluated using multivariate analysis. The relative abundance of each compound was first transformed using a centered log ratio (CLR transformation), in which the proportion of each compound was divided by the geometric mean of all compounds in the sample and then log transformed. Variation in cuticular secretion was visualized using t-distributed stochastic neighbor embedding (t-SNE) and tested using permutational multivariate analysis of variance (PERMANOVA, using the ‘adonis2’ function in the R package vegan) on a Bray–Curtis dissimilarity matrix computed from relative compound abundances, with either treatment group (callow, QR, QL) or treatment and age groups as the predictors. Because PERMANOVA can be sensitive to differences in within-group dispersion, we also tested for homogeneity of dispersion among groups (using the ‘betadisper’ function in vegan). This calculates each individual’s distance to its group centroid in multivariate cuticular lipid space. Differences in dispersion among groups were evaluated using a permutation test, and pairwise differences in mean distance to centroid were assessed using Tukey’s honestly significant difference test. To assess which compounds contributed most to group differences, we performed both linear mixed-effects models and a SIMPER analysis, which test for per-compound differences in relative or absolute abundance and partition between-group dissimilarity into per-compound contributions, respectively. These were conducted on all compounds greater than 1% of the total peak area. For the linear models (lme4), each compound was fit with treatment group as a fixed effect and colony as a random effect, and pairwise differences among groups were estimated using estimated marginal means (emmeans) with Tukey-adjusted contrasts; p-values were further FDR-corrected across all compounds and contrasts.

The total secretion and the average quantity of different chemical classes (alkanes, alkenes, acetate esters, and others) across treatments were tested using LMM with social condition, age, and their interaction as fixed effects and colony ID as a random factor, whenever needed. To compare time to death in desiccation between bees treated with solvent control and treatment, we used Kaplan Meier survival analysis with a log rank test. Finally, we compared attraction between solvent and odor control using a chi-square test. All post-hoc comparisons were obtained using a Tukey test. Statistical significance was determined at α = 0.05.

## Results

### Factors affecting death from desiccation

Time to death was significantly affected by the social condition, with QL workers surviving longer than QR workers (GLM: F_1,77_=4.55, p=0.03), and by age, with survival decreasing with age (F_1,77_=84.4, p<0.001). No interaction between social condition and age was detected (F_1, 77_=0.09, p=0.75; Figure 1A) although the effect of age is likely driven by the younger age groups. The average terminal oocyte size of workers was higher in QL than in QR at all ages (Figure S3A), and time to death was negatively correlated with oocyte size (r=-0.59, p<0.001, n=81; Figure 1B). Initial bee mass did not differ across age or social condition (Figure S3B) and was not correlated with time to death (r=-0.1, p=0.33, n=81; Figure 1C). Mass loss from the onset of the experiment until death was positively correlated with time to death (r=0.38, p=0.0004, n=81; Figure 1D). The dataset was sampled equally from the same four colonies, allowing us to test for effects of colony identity on desiccation survival, ovary activation, and body mass. None of these variables differed among colonies (time to death: F_3,77_=0.88, p=0.45; initial mass: F_3,77_=0.17, p=0.91; mass loss: F_3,77_=0.04, p=0.98; oocyte size: F_3,77_=0.15, p=0.92).

**Figure 1.**
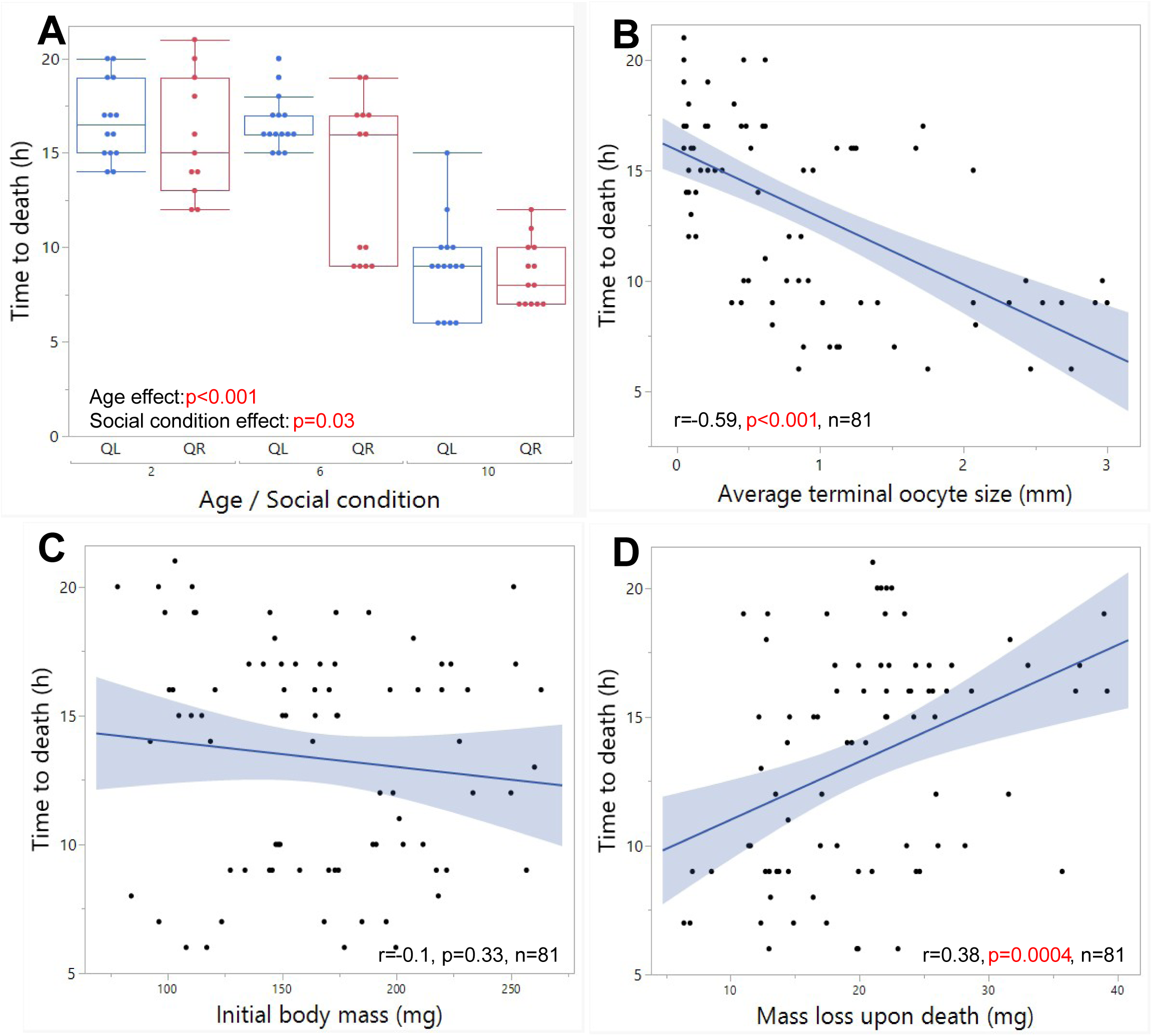
Factors affecting death from desiccation in *Bombus impatiens* workers. Queenright and queenless workers of 3 age groups (11,13,13 and 14,15,15, respectively) were placed in individual tube with silica gel at 35c and monitored hourly until death. Time to death was affected by both age and social condition (A). Time to death correlated negatively with worker ovarian activation (B) but not with initial body mass (C). Mass loss between experiment onset and death was positively correlated with time to death (D). In (A), Boxes represent the interquartile range, horizontal lines indicate medians, whiskers extend to the most extreme observations within 1.5 times the interquartile range, and points beyond the whiskers represent outliers. In (B–D), lines represent fitted regressions, and shaded areas indicate 95% confidence intervals. Data was compared using a General Linear Model and Pearson correlations

We additionally analyzed a group of newly emerged bees from the same colonies (n=15). Their ovaries were inactivated (0.05±0 mm, on average) and their initial body mass was equivalent to the other groups (140±7.4 mg, on average). However, their time to death in desiccation was significantly longer compared to the other social conditions (19.9±0.56 h, on average compared to 9-17 h in QR and QL workers (One-way ANOVA F_2,2_=596, p<0.001 followed by Tukey type post hoc p<0.001 for callows vs. both QR and QL workers).

### Cuticular lipid analysis of workers from different ages and social conditions

The cuticular lipid secretion of workers included 90 compounds (Table S2). Of these, 40 compounds were identified as alkanes and alkenes between C21 to C33 and two as acetate esters (hexacosyl acetate and tetracosyl acetate), collectively accounting for 85% of the secretion. The remaining compounds were unidentified but were not hydrocarbons, and all but two individually accounted for less than 1% of the total secretion. Subsequent analyses included only compounds that constituted at least 1% of the secretion.

Worker samples were significantly separated by age and social state. Callow bees formed a distinct group from both QR and QL workers, and the separation across social states became weaker as worker age increased. The overall PERMANOVA was highly significant (F_6,93_ = 34.47, R² = 0.69, p = 0.001), with treatment group explaining approximately 69% of the variation in cuticular lipid composition, and the seven age groups formed largely distinct clusters (Fig. 2a). All 21 pairwise PERMANOVA comparisons were significant (FDR-adjusted P ≤ 0.002) with one exception: the D10 QL vs. D10 QR contrast, which did not differ after correction (p = 0.079). The test for homogeneity of multivariate dispersion was also significant (permutational betadisper: F_6,93_ = 2.70, p = 0.02); however, Tukey HSD pairwise comparisons indicated that this result was driven by a single group pair (D2 QL vs. D10 QL; p_adj = 0.03), with the remaining 20 of 21 pairs showing homogeneous dispersion. Significant PERMANOVA differences among these 20 pairs therefore reflect genuine divergence rather than differences in dispersion. Callow workers were strongly separated from both QR and QL workers, differing significantly from every other group (pairwise PERMANOVA, all FDR-adjusted p = 0.001). The chemical distinction between QR and QL workers weakened with age: QR and QL profiles differed significantly in days 2 and 6 (FDR-adjusted p ≤ 0.005) but were statistically indistinguishable by day 10 (D10 QL vs. D10 QR, FDR-adjusted p = 0.079).

**Figure 2.**
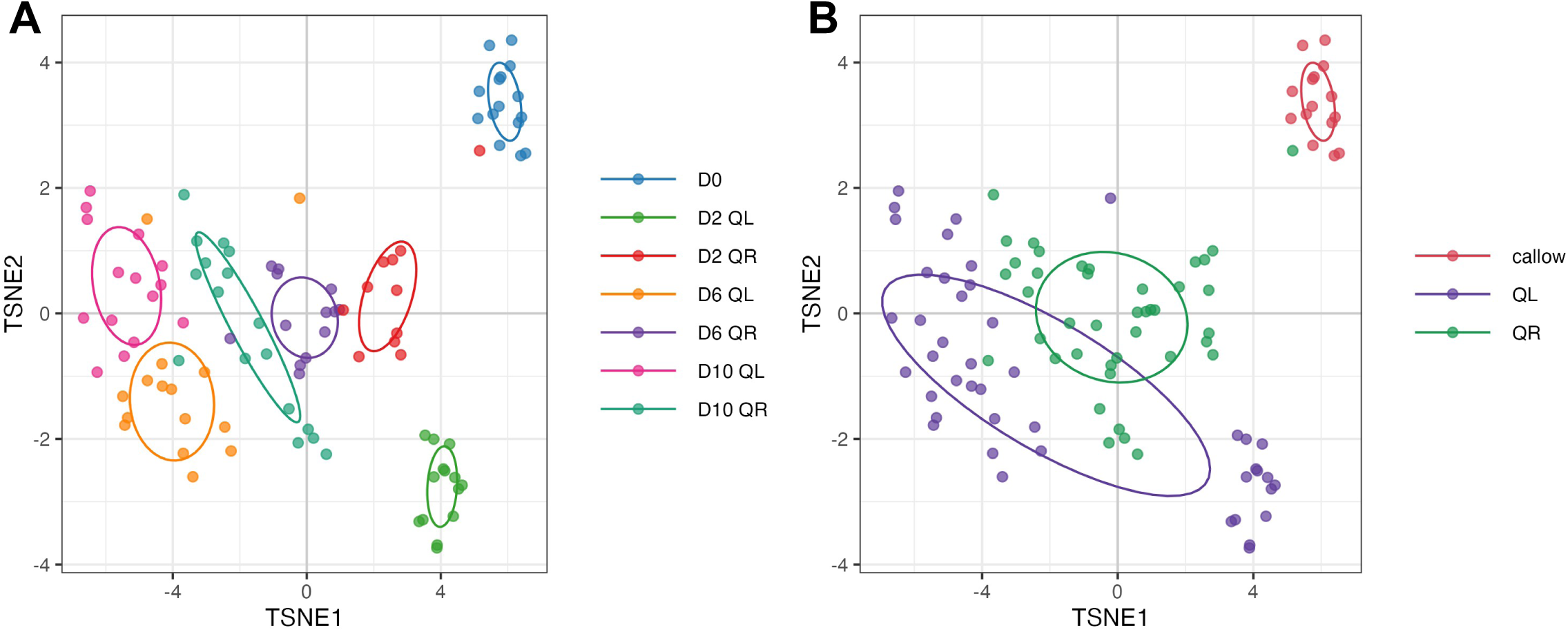
tSNE plots showing variation in cuticular lipid composition among workers of different age and social conditions (11, 13, 17 and 14,15, 15 for QR and QL, respectively). In figure A, all age and social state treatments are color-coded, whereas in figure B, only social state was color-coded. Each point represents one worker. Values are based on CLR transformed compound proportions. Ellipses represent 95% confidence regions for each group.

SIMPER analysis indicated that the top five contributors accounted for 45–55% of the average between-group dissimilarity in each pairwise comparison, with the largest and most consistent single contributor being C27 (mean contribution ∼0.06, ratio of contribution to its standard deviation 3.4–5.3 across comparisons), followed by C25, C25:1, and C27:1. Two additional alkenes, C27:1 and C23:1, were less prominent contributors but showed the highest discrimination consistency (ratios up to 6.2). Full results from the SIMPER analysis for each contrast are presented in Supplementary Table S3.

Linear mixed-effects models found similar patterns. Of the 22 abundant compounds analyzed, 18 differed significantly in at least one pairwise comparison among groups (FDR-adjusted p < 0.05), with the most consistent and largest-effect differences (7–10 significant contrasts each; Cohen’s d up to 5.6) involving alkenes: C27:1, C31:1, C29:1, and C25:1, together with the alkane C31 and the unidentified compound “comp_41”. Eleven of these compounds were significant whether analyzed as relative proportions or absolute amounts, indicating robustness to the choice of quantification scale. Full results from the linear models are presented in Supplementary Tables S4-5.

Worker total cuticular secretion was not significantly affected by age (LMM: F_1,80_.4=2.7, p=0.1) or by the interaction between age and social condition (F_1,80.4_=0.04, p=0.8). The effect of social condition on the total secretion was initially marginal (F_1,3.8_=5.8, p=0.07) but became significant (F_1,81_=4.8, p=0.03) after removing colony ID as a random factor (Figure 3A). The variance component of colony ID was insignificant (Wald p=0.58), indicating that its inclusion in the model was not justified.

**Figure 3.**
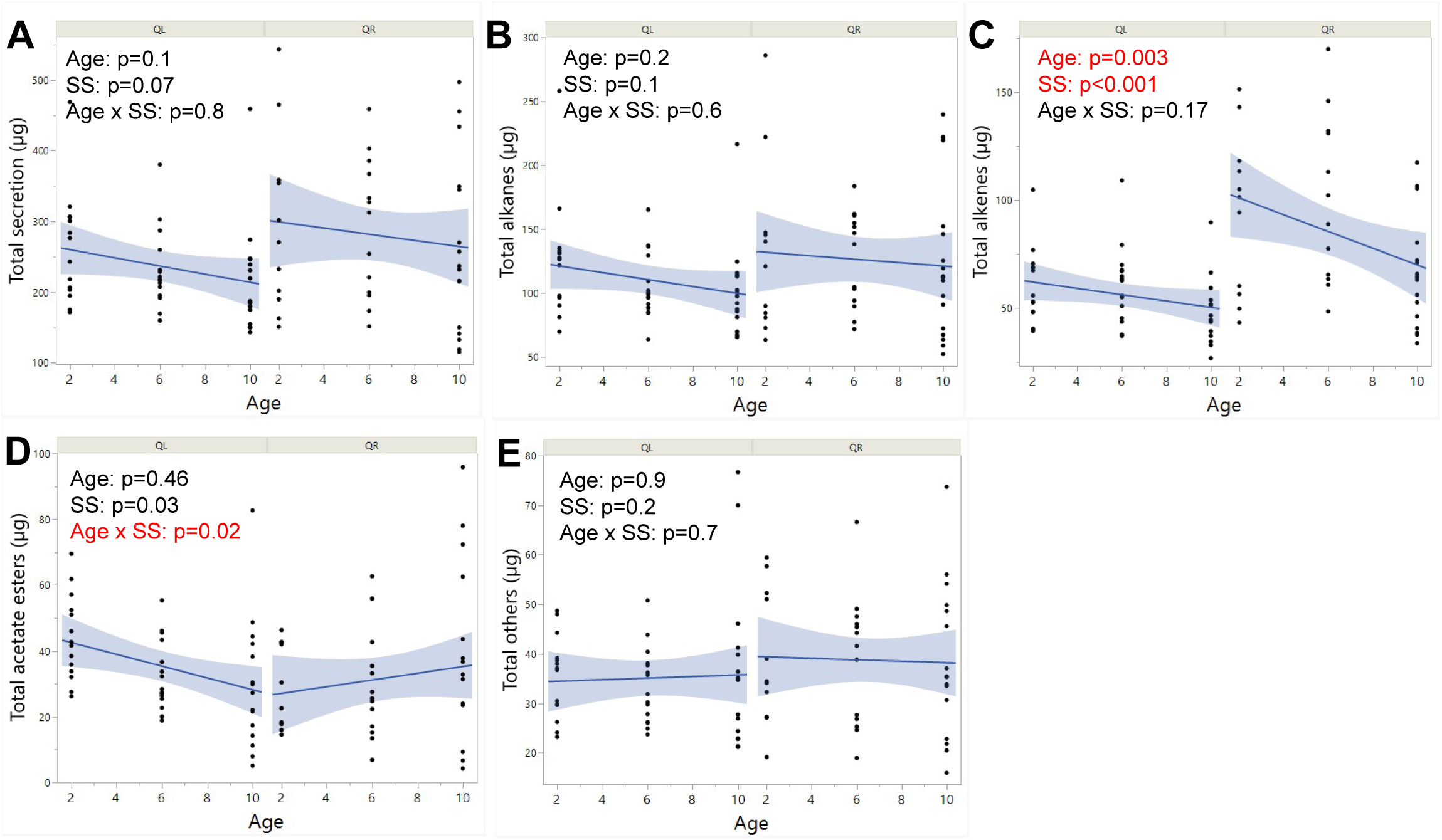
The total amount of secretion (A) alkanes (B), alkenes (C), acetate esters (D) and other compounds (E) in the cuticular lipid secretion of workers as function of age and social state (11, 13, 17 and 14,15, 15 for QR and QL, respectively). The statistical information presented in the figure refers to an LMM including age, social state (SS), and their interaction as fixed effects, with colony ID as a random effect. Points represent individual workers, lines represent fitted trends, and shaded areas indicate 95% confidence intervals.

Workers from different social states and ages did not differ in the amount of alkanes on their cuticle (age: F_1,81_=1.6, p=0.2; social state: F_1,81_=2.6, p=0.1; interaction: F_1,81_=0.18, p=0.6; Figure 3B). In contrast, the amount of alkenes was affected by both age (F_1,81_=9.05, p=0.003) and social state (F_1,81_=24.3, p<0.001), with higher amounts and a sharp decrease with age in QR workers (Figure 3C). No significant interaction was found (p=0.17). The amount of the two acetate esters was significantly affected by the interaction of age and social state (F_1,76.4_=5.04, p=0.02) but not by either factor alone (age: F_1,76.4_=0.53, p=0.46; social state: F_1,3.7_=1.44, p=0.3), with different patterns across age in each social state: in QL workers, the amount of acetate esters decreased with age, whereas in QR workers the amount increased with age (Figure 3D). We also compared the amount of unidentified compounds (neither hydrocarbons nor acetate esters, approximately 15% of the secretion) across age and social states and found no differences (age: F_1,81_=0.0002, p=0.98; social state: F_1,81_=1.65, p=0.2; interaction: F_1,81_=0.12, p=0.75; Figure 3E). Similar patterns were obtained when we compared the percentages of these chemical classes out of the total secretion (Figure S4).

We further analyzed the secretion of 15 newly emerged bees, which differed significantly from all other samples due to a limited cuticular secretion. On average, callow bees produced a total secretion of 70 µg compared to 200–400 µg in older bees. Of these, 49% were alkanes, 29% were alkenes, 7% were acetate esters, and 15% were other compounds. Thus, callow bees produced similar proportions of compound classes but about 3–4x less total secretion, likely due to immaturity of the cuticle.

### The effect of alkanes and alkenes on worker survival in desiccation

To test whether an external application of cuticular lipids can improve time to death in workers, we first applied 50 µg of ten of the most common compounds to the cuticle of newly-emerged bees (6 alkanes: C23, C24, C25, C27, C29, C31; 3 alkenes: C23:1 (7Z), C25:1 (7Z), C25:1 (9Z); and one acetate ester: tetracosyl acetate, TCA). These compounds were the most abundant on the cuticle and also contributed significantly to the separation among worker groups in the SIMPER analysis. We found no effect on time to death (C23: log rank test, χ²=1.12, p=0.28; C24: χ²=0.3, p=0.5; C25: χ²=0.3, p=0.57; C27: χ²=0.6, p=0.4; C29: χ²=0.08, p=0.7; C31: χ²=0.08, p=0.7; C23:1 (7Z): χ²=0.8, p=0.3; C25:1 (7Z): χ²=1.02, p=0.3; C25:1 (9Z): χ²=0.6, p=0.4; TCA: χ²=0.7, p=0.3; Figure S5)

We then repeated the experiment with two major changes: we applied a dose of 400 µg (x8 compared to the first round, equivalent to the amount of the total secretion on the bee cuticle) to QL bees that were 10 days old. These bees showed low survival in previous experiments and thus have greater potential for improvement in contrast to the callow bees whose survival was the highest. Treatments with five alkanes and alkenes (C23:1 (Z7), C25:1 (Z7), C27, C29, and C31) significantly extended worker lifespan under desiccation (C23:1 (7Z): χ²=4.69, p=0.03; C25:1 (Z7): χ²=5.7, p=0.01; C27: χ²=9.2, p=0.002; C29: χ²=13.02, p=0.0003; C31: χ²=5.9, p=0.03). However, a similar application with the acetate ester TCA didn’t extend worker lifespan (χ²=0.03, p=0.8; Figure 4).

**Figure 4.**
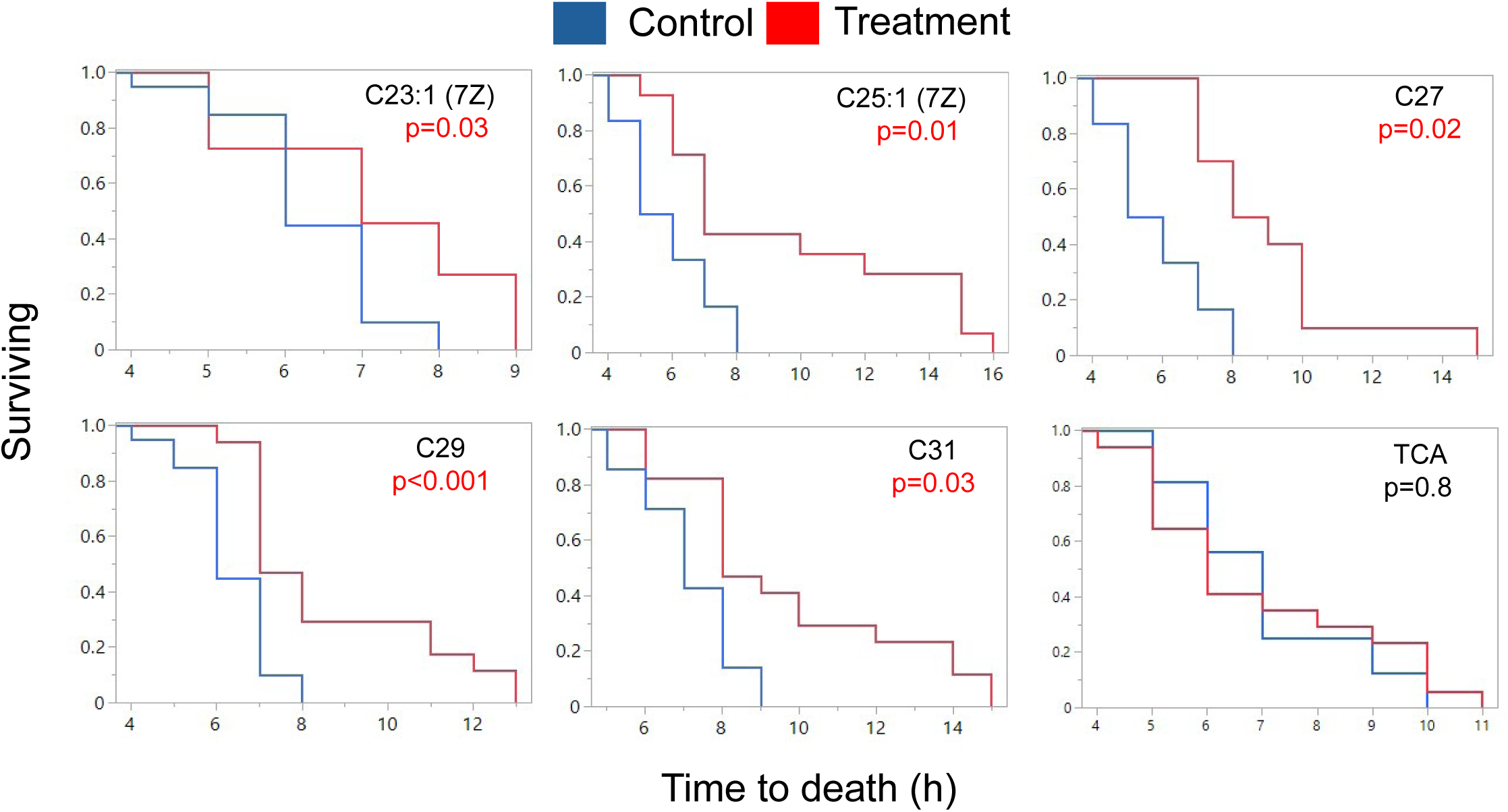
Time to death of bees treated with solvent control or treatment. 10-days old bees received a single application of 400 µg in a volume of 1 µL and were then placed under desiccation conditions. Bees were monitored hourly until death. Sample sizes (treatment/control) were 12/20, 14/6, 10/6, 17/20, 17/14, and 17/16 for C23:1, C25:1, C27, C29, C31, and TCA, respectively.

Lastly, we examined the communicative role of three selected compounds using a T maze. The response rate was 62% across all bioassays, resulting in 104 valid tests across all treatments. The response rate was higher in QR workers (70%) than in QL workers (55%). There was no preference between the control and odor treatment for either QL or QR workers when the tested odor was C25:1 or C27 (C25:1: χ² = 0.059, p = 0.8 for QL and χ² = 0.053, p = 0.8 for QR; C27: χ² = 0.06, p = 0.7 for QL and χ² = 0, p = 1 for QR). In contrast, QR workers showed significant attraction to TCA over the control (χ² = 6.3, p = 0.01), whereas QL workers showed no significant preference between TCA and the control (χ² = 2.2, p = 0.1). QL workers further selected the control twice as often as TCA (11 vs. 5), suggesting a slight trend toward TCA repellency. Overall, attraction depended on the social state and was observed only in QR workers in response to TCA (Figure 5).

**Figure 5.**
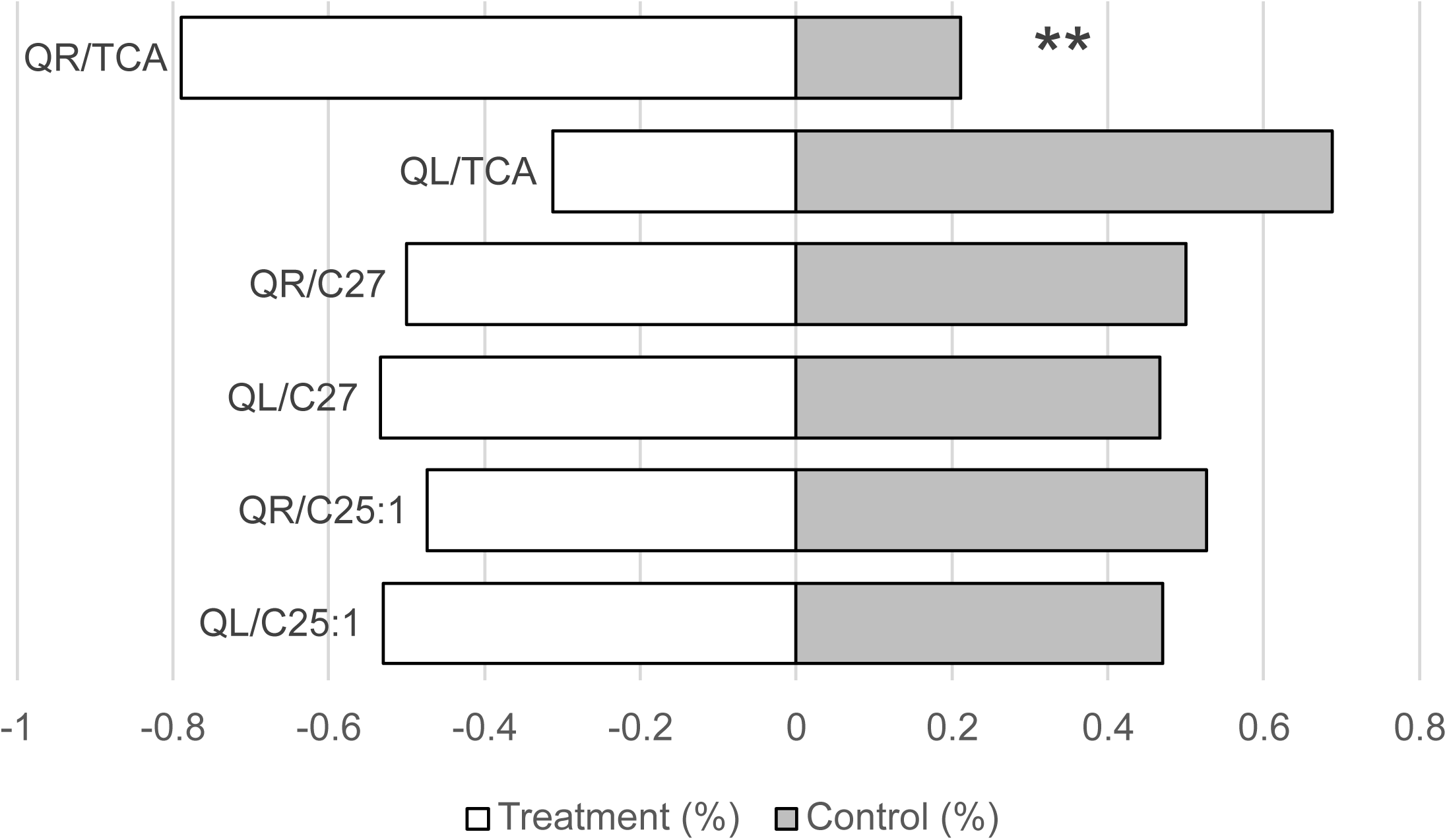
Attraction of queenright (QR) and queenless (QL) workers to selected cuticular compounds, tetracosyl acetate (TCA), heptacosane (C27), and pentacosene (C25:1 (Z7)). Workers were given 10 minutes in a T maze to choose between a solvent control (hexane) and an odor treatment (400 µg/1 µL). Asterisks denote significance at p = 0.01. Sample sizes (QL/QR) include 28/26, 28/26 and 32/28 for C25:1, C27 and TCA, respectively.

## Discussion

Our study demonstrates a trade-off between desiccation resistance and producing communicative signals in bees. This trade-off is not simply driven by differences in the quantities of protective and signaling compounds, but rather by age-dependent shifts in their relative proportions, particularly in the presence of the queen. The insect cuticular lipid layer serves as the primary hydrophobic barrier against water loss (Gibbs, 1998, Blomquist and Ginzel, 2021) and it varies with internal factors such as age and reproductive status. Worker survival under desiccation stress decreased with age and ovarian activation (Lyons et al., 2014). Theoretically, because QR workers activate their ovaries more slowly than QL workers, they were expected to survive longer under desiccation stress. However, QR workers did not survive longer; instead, QL workers of the same age showed higher survival overall. We hypothesized that desiccation survival in QR workers is impaired by a trade-off with cuticular signals used for communication as they age. Unlike QL workers, which prioritize direct reproduction, QR workers live in a complex social environment in which direct reproduction is suppressed and communication with the queen and nestmates is intensified (Amsalem et al., 2015). We further explored this explanation by testing the role of selected cuticular compounds in desiccation resistance and communication, and found a striking dichotomy: several hydrocarbons, but not the acetate ester we examined, improved desiccation resistance, whereas attraction was demonstrated only for the acetate ester and was specific to the QR social state. Together, these findings suggest a trade-off between the protective and communicative functions of cuticular secretions, reflected in age-dependent shifts in their composition.

Analysis of the cuticular secretion revealed that desiccation survival was not driven by the total amount of secretion. Instead, the differences in survival were more likely driven by secretion composition, specifically age-dependent decrease in the amounts of alkenes and age-dependent increase in the amounts of acetate esters in QR compared to QL workers were associated with lower survival in desiccation. Both alkanes and alkenes contributed to desiccation resistance, whereas the acetate ester TCA provided no such benefit, as further confirmed by survival assays following topical application of compounds from these groups. Because cuticular lipids form a compositional mixture, changes in the relative abundance of individual compounds are inherently interdependent: an increase in the relative proportion of some compounds necessarily comes at the expense of others. In other words, a higher proportion of hydrocarbons on the cuticle provides greater protection against desiccation, but some of these hydrocarbons, particularly alkenes, are reduced in QR workers as they age, potentially reflecting the need to coat the cuticle with additional compounds (acetate esters) that are essential for communication. Confounding factors such as colony ID, group size, and housing conditions cannot be fully disentangled from social condition in the present study. An experimental design in which workers from the same colonies are split between the two treatments would allow these potential effects to be disentangled. However, differences between QR and QL workers are unlikely to be driven primarily by these factors, as previous studies have emphasized the dominant contribution of reproductive state to chemical differentiation (Derstine et al., 2023, Derstine et al., 2021, Amsalem et al., 2014b, Amsalem et al., 2014a, Amsalem and Hefetz, 2011, Amsalem et al., 2009).

Between alkanes and alkenes, greater protection against water loss is likely provided by alkanes, which contribute more strongly to tight packing and waterproofing because they are linear and fully saturated. In contrast, unsaturated compounds such as alkenes, although still contributing to protection against water loss compared to other cuticular compounds, disrupt tight molecular packing, lower the melting point, and increase permeability of the lipid layer (Leonhardt et al., 2016, Hefetz, 2026). Nevertheless, topical application of both alkanes and alkenes increased survival under desiccating conditions. The positive effect of alkenes, despite their expected increase in cuticular permeability, may result from increasing the overall quantity of hydrophobic lipids on the cuticle or from forming an additional surface barrier rather than altering the composition of the endogenous lipid layer. The total cuticular secretion is likely governed by a zero-sum trade-off between the production of various compounds. Thus, if acetate esters are added for the purpose of communication while minimizing the impact on desiccation resistance, reducing alkenes rather than alkanes would be expected. Consistent with this idea, alkane amounts remained relatively stable across ages and social states, whereas alkenes and acetate esters varied substantially. At the biosynthetic level, all of these compounds are derived from the same fatty acid precursors (Derstine et al., 2023). Thus, the observed trade-off could result from increased allocation of long chain fatty acyl intermediates toward the production of fatty alcohols and acetate esters.

These relationships are further complicated by the physiological changes that occur during maturation. We observed a reduction in desiccation resistance with age, consistent with patterns reported across insect taxa, in which younger adults exhibit greater desiccation tolerance than older individuals (Lyons et al., 2014). This decline aligned with a reduction in total cuticular secretion over time, with a notable exception in callow bees, which possessed lower amounts of secretion yet showed the highest survival, likely because of their limited mobility and potentially lower metabolic rate. In the first round of topical application, we used callow bees because their cuticular secretion is still immature, but this approach proved uninformative because callow bees already maximized their survival under extreme desiccation conditions and their survival could not be further improved. In the second round, we used 10-day-old QL workers, a design that proved more effective given the significantly impaired survival of these older bees under desiccation stress. Differences in metabolic rate are unlikely to explain the greater desiccation survival of QL workers. QL workers establish dominance hierarchies and engage in more aggressive interactions than QR workers (Amsalem et al., 2013a, Amsalem and Hefetz, 2011, Amsalem and Hefetz, 2010), which should increase activity and metabolic water loss and thus, if anything, reduce rather than enhance desiccation survival.

These chemical differences between QL and QR workers likely reflect two distinct physiological and ecological priorities. Production of acetate esters in QR workers with age is likely important for social interactions, whereas QL workers prioritize reproduction and survival and maintain only minimal social interactions with nestmates, most of which are aggressive. Although the precise function of these acetate esters remains unknown, we confirmed their likely communicative role in a T maze assay showing selective attraction to TCA, but not to hydrocarbons, and only by QR workers. TCA was previously identified on the cuticle of *B. impatiens* queens and is significantly more abundant in young egg laying queens (Orlova et al., 2020). It was also identified in young larvae (unpublished data, Derstine et al.), which require intensive care. Its abundance in young colonies could explain why QR, but not QL workers are attracted to it. Across social Hymenoptera, various acetates function as alarm pheromones (Hunt et al., 2003, Du et al., 2021), recognition signals (Breed, 1998), and highly specific triggers for hygienic brood care (Noël et al., 2025). Compounds such as acetate esters possess a degree of polarity that may facilitate interactions with olfactory receptor neurons, potentially increasing detectability and signal specificity (Du and Chen, 2026, Lhomme et al., 2015). Finally, it should be noted that volatility is unlikely to explain the attraction to TCA, since the only positive attraction observed in this study was toward TCA, the heaviest compound among those tested.

The age-dependent trade-off between acetate esters and alkenes in QR workers is likely possible only because these workers function mostly within the buffered, humid microclimate of the nest and do not activate their ovaries in young colonies. This protected environment may relax the selective pressure for individual waterproofing, allowing more complex chemical communication to evolve (Perez and Aron, 2020). In contrast, conditions outside the nest leave QL workers functionally isolated and more vulnerable to unbuffered environmental exposure. This, together with a reduced need for communication and a greater prioritization of reproduction and survival, likely favors investment in a heavier and more tightly packed hydrocarbon layer.

## Conclusion

In conclusion, desiccation tolerance in *B. impatiens* workers is a dynamic trait regulated by the social environment. QR workers appear to trade investment in alkenes for acetate esters as they age, effectively reducing their resistance to desiccation while increasing the production of communicative compounds associated with nestmates, the queen, and likely also the brood. Understanding how insects resolve such trade-offs is central to explaining the evolution and plasticity of cuticular lipid profiles and may provide broader insight into the selective forces shaping social insect societies.

## Acknowledgments

We would like to thank Prof. Henry Chung from Michigan State University for inspiring this study and providing information about the apparatus used to measure desiccation survival.

## Author Contribution

SAMH, ND and EA conceived the ideas and designed methodology. SAMH collected and analyzed the data and wrote the first draft. ND conducted the chemical analysis. EA analyzed and visualized the data and wrote the manuscript together with SAMH and ND. All authors contributed critically to the drafts and gave final approval for publication.

## Data availability statement

The data of this study is provided in Tables S6-9 in the supplementary materials.

