## Supplemental figures for "A trade-off between desiccation resistance and social communication in a social bee"

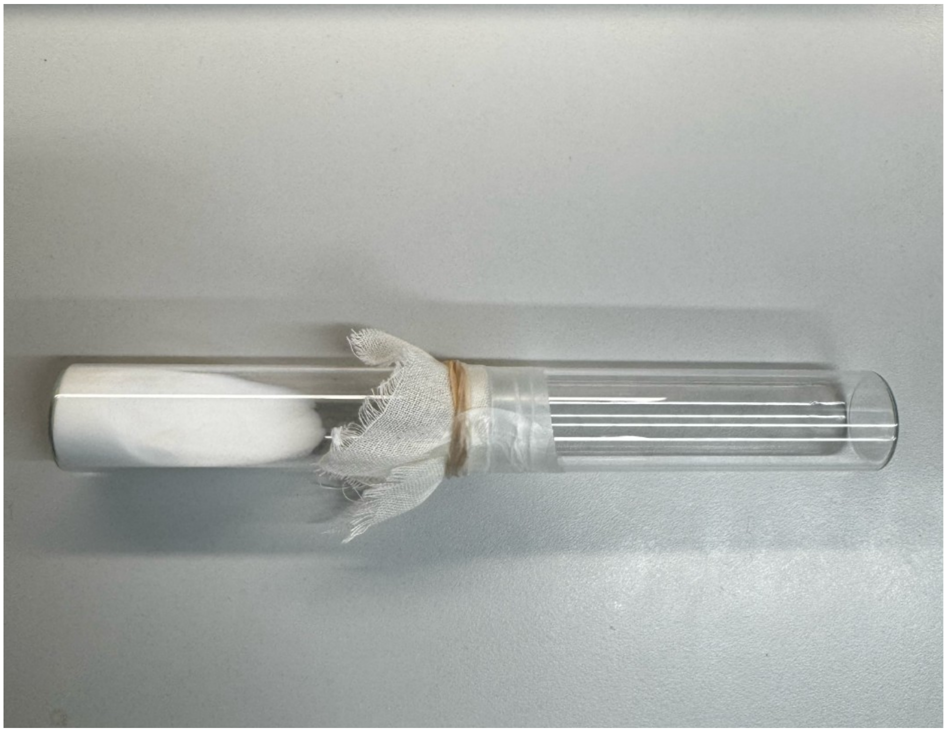

**Figure S1**. The apparatus used for desiccation assays. The left arm was filled with silica gel, whereas the bee was placed in the right arm. The two arms were separated by mesh.

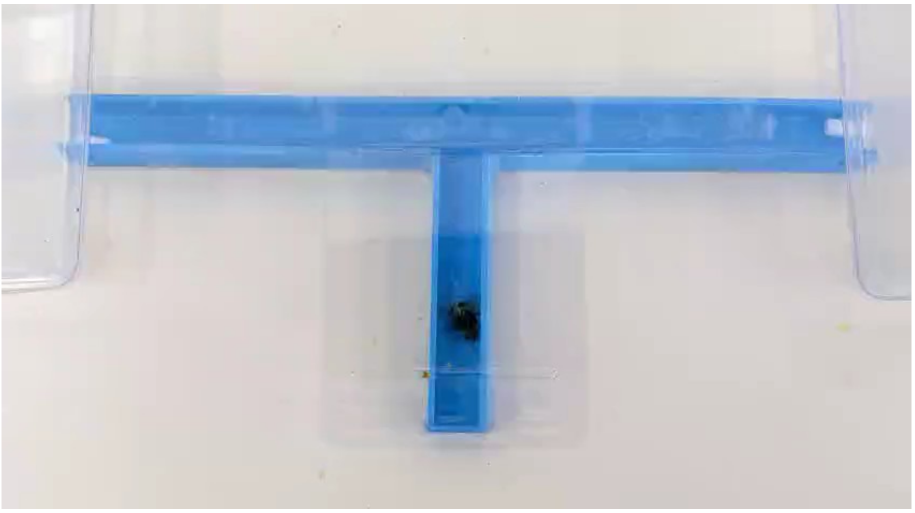

**Figure S2**. A two-choice olfactometer bioassay containing one worker choosing between a solvent control and a treatment odor.

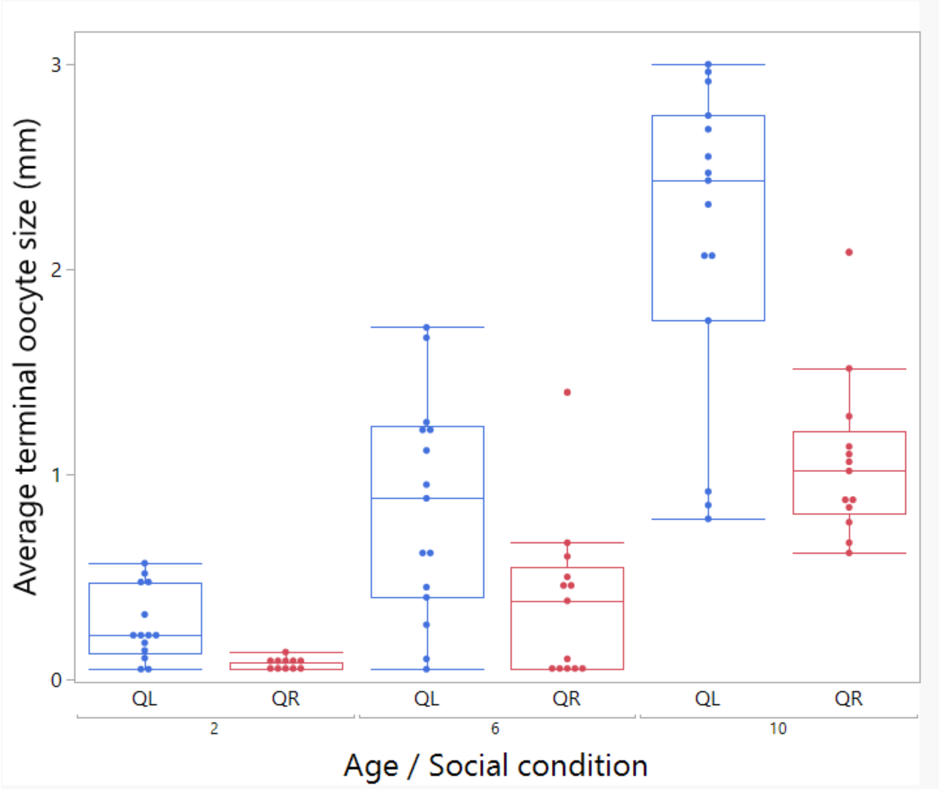


**Figure S3A**. The average terminal oocyte size was significantly affected by age ($F_{1,77}$ = 115.8, p < 0.001), social condition ($F_{1,76.4}$ = 30.8, p < 0.001), and their interaction ($F_{1,74.6}$ = 11.5, p < 0.001), with QL workers showing higher ovarian activation overall and activating their ovaries faster with age.

Colony ID was included as a random factor, but its contribution was not significant. Boxes represent the interquartile range, horizontal lines indicate medians, whiskers extend to the most extreme observations within 1.5 times the interquartile range, and points beyond the whiskers represent outliers. Sample sizes include 11,13,13 and 14,15,15 workers for QR and QL in age groups 2,6 and 10, respectively.

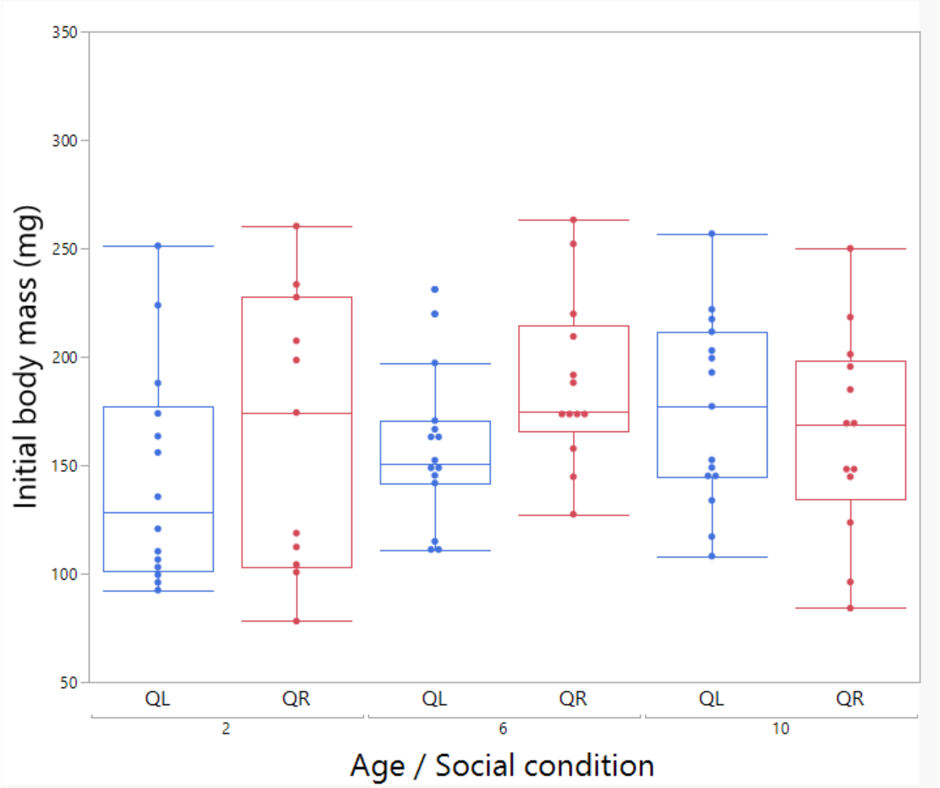


**Figure S3B**. The initial body mass of workers was not significantly affected by age ($F_{1,42.1} = 0.36$, p = 0.55), social condition ($F_{1,66.6} = 1.61$, p = 0.2), or their interaction ($F_{1,76.1} = 1.82$, p = 0.18). Colony ID contribution was insignificant. Boxes represent the interquartile range, horizontal lines indicate medians, whiskers extend to the most extreme observations within 1.5 times the interquartile range, and points beyond the whiskers represent outliers. Sample sizes include 11,13,13 and 14,15,15 workers for QR and QL in age groups 2,6 and 10, respectively.

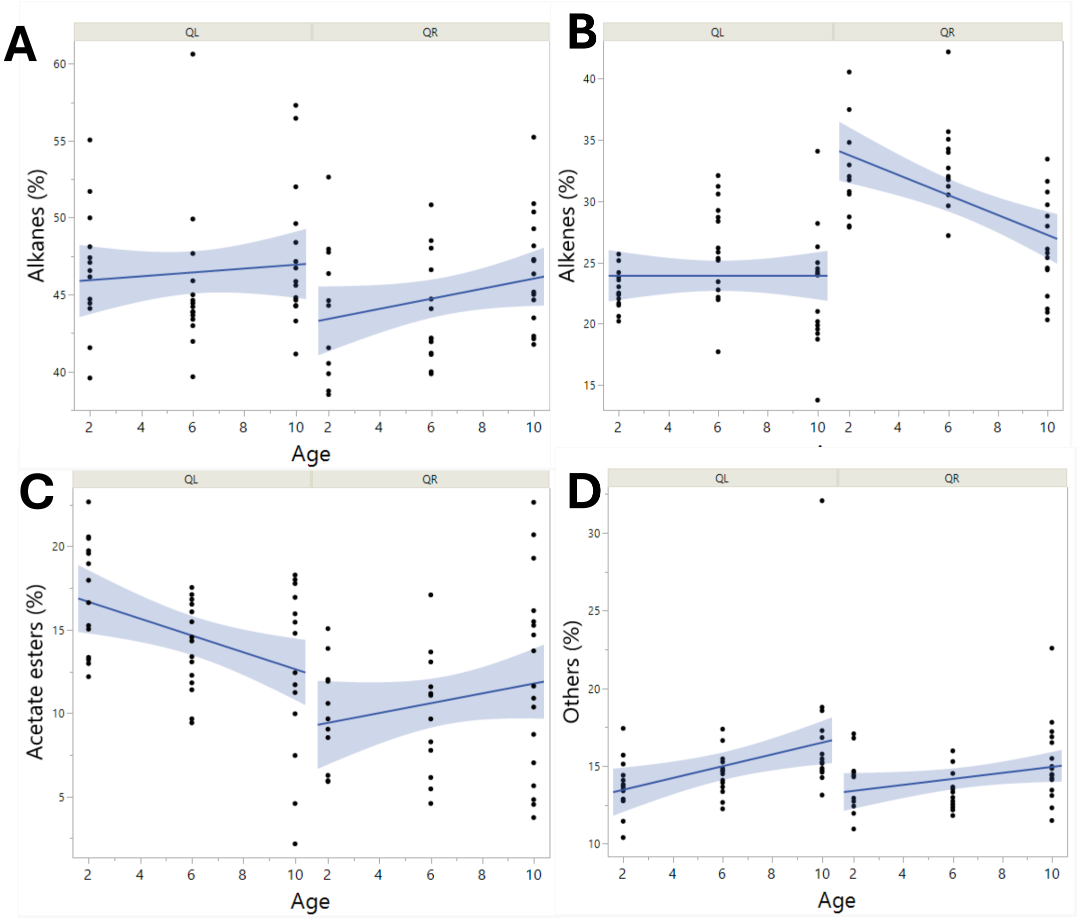


**Figure S4**. The percentages of alkanes, alkenes, acetate esters and other compounds in the cuticular lipid secretion of workers as function of age and social state. Points represent individual workers, lines represent fitted trends, and shaded areas indicate 95% confidence intervals. Sample sizes include 11, 13, 17 and 14,15, 15 workers for QR and QL, respectively.

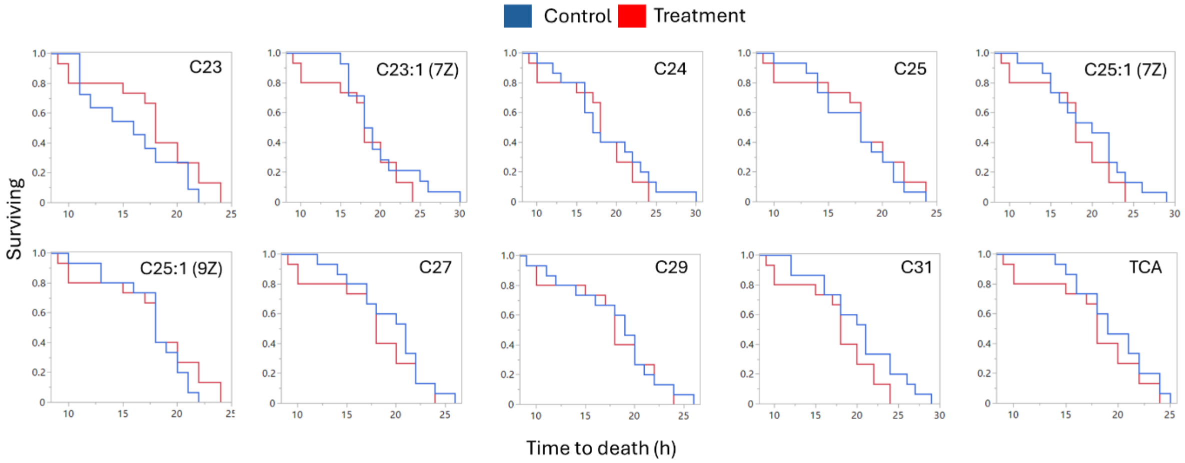


**Figure S5**. Time to death of bees treated with solvent control or treatment. Newly emerged bees received a single application of 50 µg and were then placed under desiccation conditions. Bees were monitored hourly until death. n=11–15 per treatment. None of the comparisons were significant ($p>0.05$). Sample sizes (treatment/control) were 11/15 (C23), 14/15 (C23:1), and 15/15 for the remaining 8 compounds.
